# Structural insights into secretory immunoglobulin A and its interaction with a pneumococcal adhesin

**DOI:** 10.1101/2020.02.11.943233

**Authors:** Yuxin Wang, Guopeng Wang, Yaxin Li, Hao Shen, Huarui Chu, Ning Gao, Junyu Xiao

**Author notes:** These authors contributed equally to this work (Yuxin Wang, Guopeng Wang, Yaxin Li). Correspondence and requests for materials should be addressed to J.X.

## Abstract

Secretory Immunoglobulin A (SIgA) is the most abundant antibody at the mucosal surface. SIgA possesses two additional subunits besides IgA: the joining chain (J-chain) and secretory component (SC). SC is the ectodomain of the polymeric immunoglobulin receptor (pIgR), which functions to transport IgA to the mucosa. The underlying mechanism of how the J-chain and pIgR/SC facilitates the assembly and secretion of SIgA remains to be understood. During the infection of *Streptococcus pneumoniae*, a pneumococcal adhesin SpsA hijacks SIgA and unliganded pIgR/SC to evade host defense and gain entry to human cells. How SpsA specifically targets SIgA and pIgR/SC also remains unclear. Here we report a cryo-electron microscopy structure of the Fc region of human IgA1 (Fcα) in complex with J-chain and SC (Fcα-J-SC), which reveals the organization principle of SIgA. We also present the structure of Fcα-J-SC in complex with SpsA, which uncovers the specific interaction between SpsA and human pIgR/SC. These results advance the molecular understanding of SIgA and shed light on the pathogenesis of *S. pneumoniae*.

## Introduction

The mucous membrane covers ~400 m^2^ surface of internal organs in the human body. Immunoglobulin A (IgA) is the most predominant antibody present at the mucosa ^1^. In contrast to IgA in serum that is mostly monomeric, mucosal IgA are mainly present as dimers (dIgA), in which two IgA molecules are linked together by another protein designated the joining chain (J-chain) ^2^. The J-chain is also present in the IgM pentamer (pIgM) and facilitates its assembly ^3^. The heavy chains of IgA and IgM contain unique C-terminal extensions known as the tailpieces, which are essential for their oligomerization and covalent linkage to the J-chain ^4–6^. Furthermore, an additional polypeptide called the secretory component (SC) is present in mucosal IgA and IgM, and such IgA and IgM complexes are often referred to as secretory IgA and IgM (SIgA and SIgM). SC is the ectodomain of the polymeric immunoglobulin receptor (pIgR), which functions to transport dIgA and pIgM through the mucosal epithelial cells ^7–10^. SIgA forms a critical first line of defense against pathogens at the mucosal surface, and also likely plays an important role in regulating the homeostasis of microbiota ^11,12^. SIgA in breast milk is important for protecting the newborn babies until their own immune systems have developed. Despite the fact that the composition of SIgA and certain details of its assembly process have long been established, the three-dimensional structure of SIgA has remained elusive.

Due to the critical function of SIgA in immune defenses, various pathogens have developed strategies to disrupt its function. *Streptococcus pneumoniae*, also known as pneumococcus, is a Gram-positive bacterium that causes millions of deaths worldwide ^13,14^. It is an opportunistic pathogen residing in the upper respiratory tract of many people especially young children. In individuals with a weak immune system, the bacterium can invade a wide range of organs including the brain, causing severe diseases such as pneumonia, sepsis, and meningitis. *<u>S. p</u>neumoniae* <u>S</u>Ig<u>A</u> binding protein (SpsA; also known as CbpA, PspC) is a pneumococcal adhesin that binds to SIgA ^15^. The binding is mediated by SC, and may impair the bacterial clearance function of SIgA. Furthermore, SpsA also interacts with unliganded SC and pIgR, and the interaction with pIgR may enhance bacterial adherence and importantly, facilitate its cellular invasion ^16^. How SpsA selectively recognizes human pIgR/SC remains to be characterized.

Here we report a cryo-electron microscopy (cryo-EM) structure of the human Fcα-J-SC complex at 3.2 Å resolution. Comparison of this structure with that of Fcμ-J-SC ^17^ reveals a more complete structure of the J-chain and distinctive features for the interactions between Fcα, J-chain, and SC. We also investigated the interaction between SIgA and the *S. pneumoniae* adhesin SpsA, and determined a cryo-EM structure of human Fcα-J-SC in complex with the N-terminal domain of SpsA, which shows how human pIgR/SC is specifically exploited by SpsA to promote *S. pneumoniae* pathogenesis.

## Results

### Overall structure of the Fcα-J-SC complex

We co-expressed human IgA1-Fc (Fcα) with J-chain in HEK293F cells and isolated the complex containing the dimeric Fcα. SC was individually expressed and purified, and then incubated with the Fcα-J sample to form the Fcα-J-SC tripartite complex. We then determined its structure at 3.2 Å resolution (as judged by the FSC 0.143 criterion) by the single particle cryo-EM method (Fig. 1 and Supplementary information, Fig. S1). Most regions of the EM map exhibited atomic resolutions, allowing unambiguous structural assignment and analyses (Supplementary information, Fig. S1e and Fig. S2). The statistics for cryo-EM data collection and processing, as well as structural refinement and validation, are summarized in Supplementary information, Table S1.

**Figure 1.**
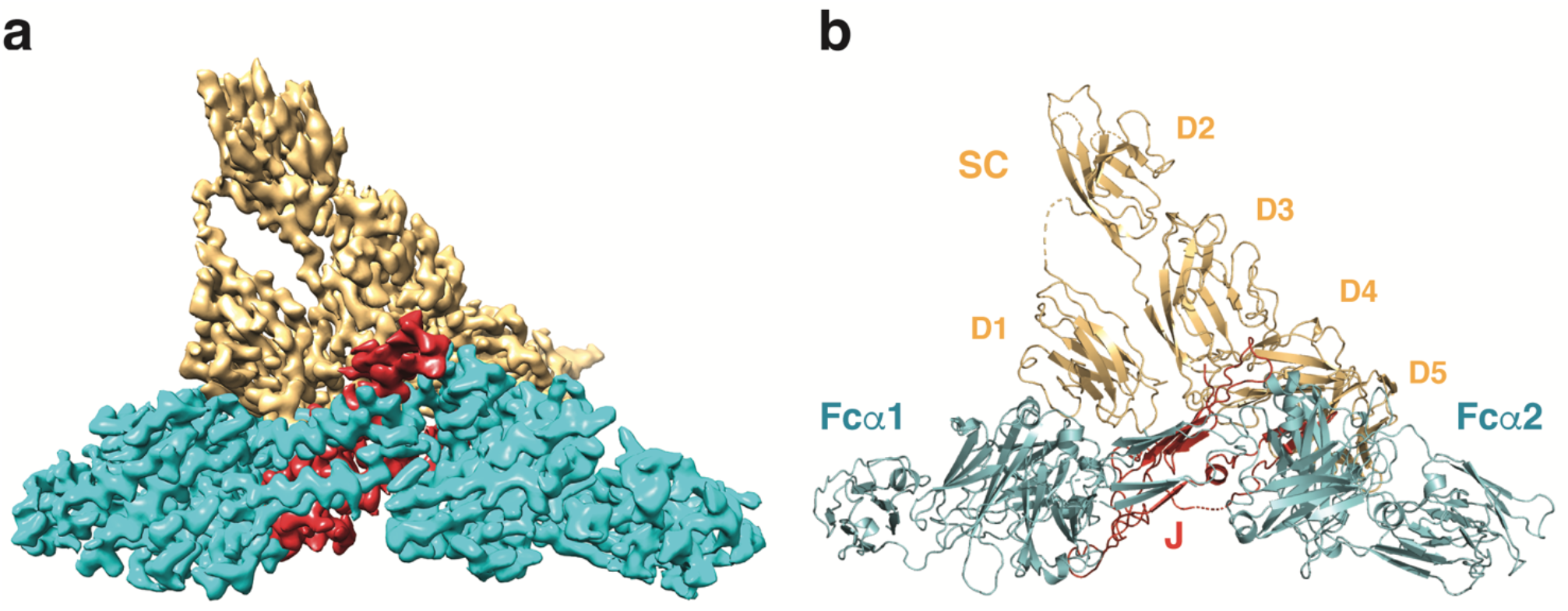
Cryo-EM structure of the human Fcα-J-SC complex. **a.** The cryo-EM density map of Fcα-J-SC reconstructed at 3.15 Å resolution. The regions corresponding to Fcα, J-chain, and SC are shown in cyan, red, and gold, respectively. The same color scheme is used in all figures unless otherwise indicated. **b.** The structural model of Fcα-J-SC. The five immunoglobulin-like domains in SC are indicated as D1-D5.

The two IgA molecules in SIgA are linked via Cys471-mediated disulfide bonds between each other and to the J-chain. Earlier EM analyses show that dIgA displays a double-Y-like shape ^18^. Solution scattering studies suggest that the two Fc regions do not dock to each other in a straight manner in the IgA dimer, but adopt a slightly bent end-to-end arrangement ^19^. Consistently, our structure shows that the Fcα dimer has a boomerang-like shape (Fig. 2a) and resembles a portion of the pIgM structure we determined recently ^17^ (Fig. 2b). Each tailpiece of Fcα contains a β-strand, like that of Fcμ in IgM, and the four tailpiece strands bundle together to mediate the interactions between the two Fcα molecules. The dimeric structure is further stabilized by the J-chain.

**Figure 2.**
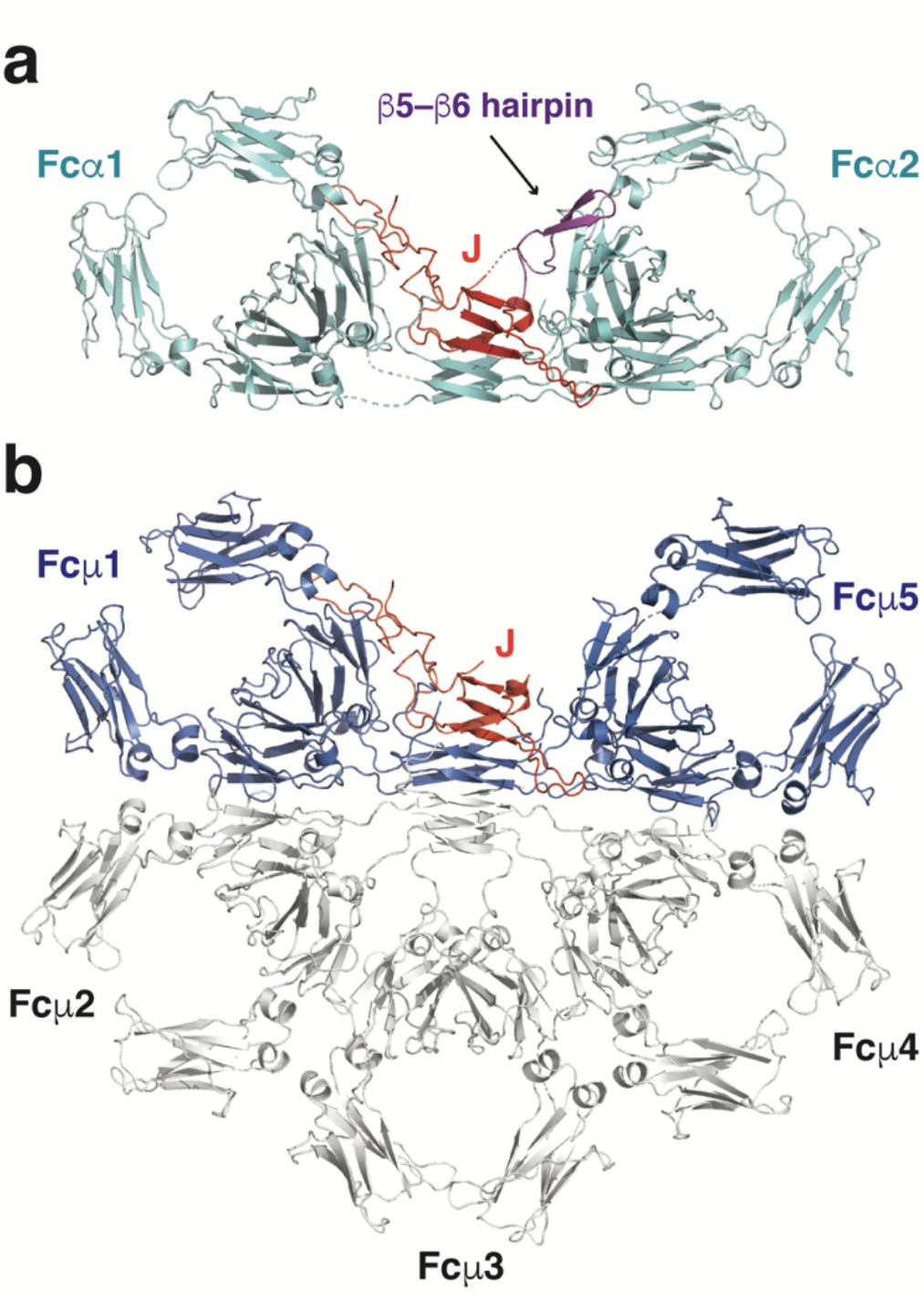
Structure of dIgA core and its comparison with pIgM. **a.** Overall structure of the dimeric Fcα in complex with the J-chain. The β5-β6 hairpin of the J-chain that is disordered in the Fcμ-J structure is highlighted in magenta. **b.** Structure of the pentameric Fcμ in complex with the J-chain. Fcμ1 and Fcμ5 are shown in blue, whereas Fcμ2-4 are shown in white.

### Interaction between IgA and the J-chain

Compared to the J-chain in the Fcμ-J complex, a more complete structure of the J-chain is present in Fcα-J, due to more extensive interactions between the J-chain and the Fcα dimer. Residues 70-92, disordered in the Fcμ-J-SC structure, form a β-hairpin (β5-β6 hairpin) that interacts with Fcα2 (Fig. 2a and Fig. 3a).

**Figure 3.**
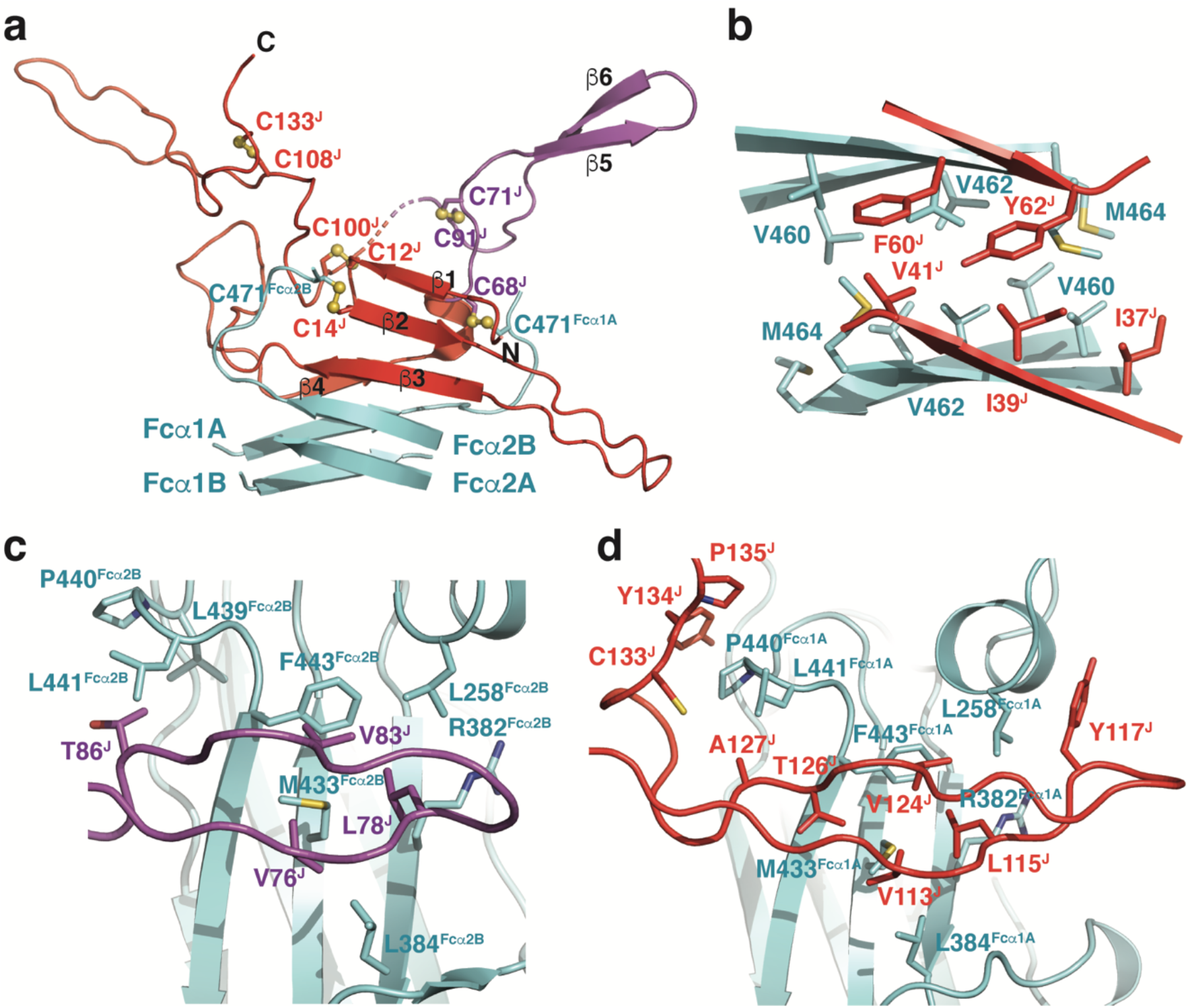
The interactions between the J-chain and Fcα. **a.** Overall structure of the J-chain. The sulfur atoms in the Cys residues that form disulfide bonds are depicted in orange spheres. The β-strands in the J-chain and Fcα tailpieces are indicated. **b.** Interactions between two J-chain strands and the Fcα tailpieces. **c.** Interactions between the β5-β6 hairpin of the J-chain and the Cα2-Cα3 junction of Fcα2β. **d.** Interactions between the C-terminal hairpin of the J-chain and the Cα2-Cα3 junction of Fcα1A.

Three intrachain disulfide bridges are present within the J-chain (Cys12^J^-Cys100^J^, Cys71^J^-Cys91^J^, and Cys108^J^-Cys133^J^; superscript J indicates J-chain residues), consistent with previous analyses ^20,21^. Cys14^J^ and Cys68^J^ form a disulfide bond with Cys471^Fcα2B^ and Cys471^Fcα1A^, respectively.

The central region of the J-chain contains four β-strands (β1-β4) that interact with the Fcα tailpieces (Fig. 3a). Strands β1-β3 pack onto the two tailpiece strands of Fcα2 to assemble into a β-sheet, with hydrogen bonds formed between main chain atoms of adjacent strands; whereas β4 packs onto the tailpiece strands of Fcα1. Robust hydrophobic interactions are present between the two β-sheets, mediated by Fcα residues Val460, Val462, Met464 and J-chain residues Ile37^J^, Ile39^J^, Val41^J^, Phe60^J^, Tyr62^J^ (Fig. 3b). The β2-β3 loop, β3-β4 loop, β5-β6 hairpin, and the long C-terminal hairpin of the J-chain function as four lassos that assist to further interact with Fcα1 and Fcα2. The β2-β3 loop interacts with the Cα3-tailpiece junction of Fcα2B (Supplementary information, Fig. S3a). Ile21^J^ and Val33^J^, together with Ile5^J^ and Leu7^J^ in the N-terminal region of the J-chain, form a hydrophobic pocket to accommodate Leu451^Fcα2B^. Asp31^J^ forms a salt bridge with Arg450^Fcα2B^. The β3-β4 loop contacts the Cα3-tailpiece junction of Fcα1A (Supplementary information, Fig. S3b). Arg46^J^ forms an ion pair with Asp449^Fcα1A^. Leu56^J^ packs on Leu451^Fcα1A^. More prominent interactions with Fcα are mediated by the β5-β6 and C-terminal hairpins of the J-chain. The β5-β6 hairpin forms extensive interactions with the Cα2-Cα3 junction of Fcα2B via two hydrophobic centers (Fig. 3c). The first is formed between J-chain residues Val76^J^, Leu78^J^, Val83^J^, and Fcα2B residues Leu258^Fcα2B^, Arg382^Fcα2B^ aliphatic side chain, Leu384^Fcα2B^, Met433^Fcα2B^, Phe443^Fcα2B^. The second hydrophobic center focuses on Thr86^J^, which is surrounded by Leu439^Fcα2B^, Pro440^Fcα2B^, and Leu441^Fcα2B^. Several hydrogen bonds are also formed between the J-chain and Fcα2B at this region, involving Asp79^J^, Thr86^J^, Asn89^Fcα2B^, Val349^Fcα2B^, Glu389^Fcα2B^, and Phe443^Fcα2B^. The C-terminal hairpin targets almost the same region in Fcα1A (Fig. 3d). Val113^J^, Leu115^J^, Tyr117^J^, Val124^J^, and Thr126^J^ mingle with Leu258^Fcα1A^, Arg382^Fcα1A^, Leu384^Fcα1A^, Met433^Fcα1A^, and Phe443 ^Fcα1A^. Ala127^J^, Cys133^J^, Tyr134^J^, and Pro135^J^ encircle Pro440^Fcα1A^ and Leu441^Fcα1A^. The way that the C-terminal hairpin is attached to Fcα1A highly resembles how it binds to Fcμ1 in IgM (Fig. 2a, 2b).

### Interaction between dIgA and SC

The interaction between dIgA and pIgR/SC has been extensively studied by biochemical and biophysical studies ^22–28^. pIgR/SC forms a bidentate interaction with dIgA, with both its D1 and D5 domain involved. The D1 domain of pIgR/SC binds to Fcα-J using its three CDR (complementarity determining regions) loops, and the molecular interactions are in many ways similar to the interactions seen in the Fcμ-J-SC complex ^17^. CDR1 mainly contacts the J-chain (Fig. 4a). Val29^SC^ (superscript SC indicates pIgR/SC residues) is positioned in a pocket formed by J-chain residues Arg105^J^, Asn106^J^, and A132^J^ to mediate hydrophobic/van der Waals interactions. Asn30^SC^ coordinates Arg105^J^. Arg31^SC^ interacts with Asp136^J^. His32^SC^ packs against Tyr134^J^. Besides these interactions with the J-chain, pIgR/SC also directly interacts with IgA at several places (Fig. 4b). For example, Arg34^SC^ in CDR1 forms a salt bridge with Glu363^Fcα1B^, the aliphatic side chain of which also packs on Tyr55^SC^ in CDR2. The main chain carbonyl of Cys46^SC^ forms a hydrogen bond with Asn362^Fcα1B^. Glu53^SC^ interacts with Arg346^Fcα1A^. Gly54^SC^ is covered by Phe345 ^Fcα1A^ and Thr408 ^Fcα1A^. Arg99^SC^ and Leu101^SC^ in CDR3 encloses Tyr472^Fcα2B^, the terminal residue of Fcα2B, together with Arg105^J^ (Fig. 4a). Two SC mutants, V29N/R31S and R99N/L101T, which display greatly reduced interactions with the Fcμ-J complex ^17^, also failed to bind Fcα-J (Fig. 4c).

**Figure 4.**
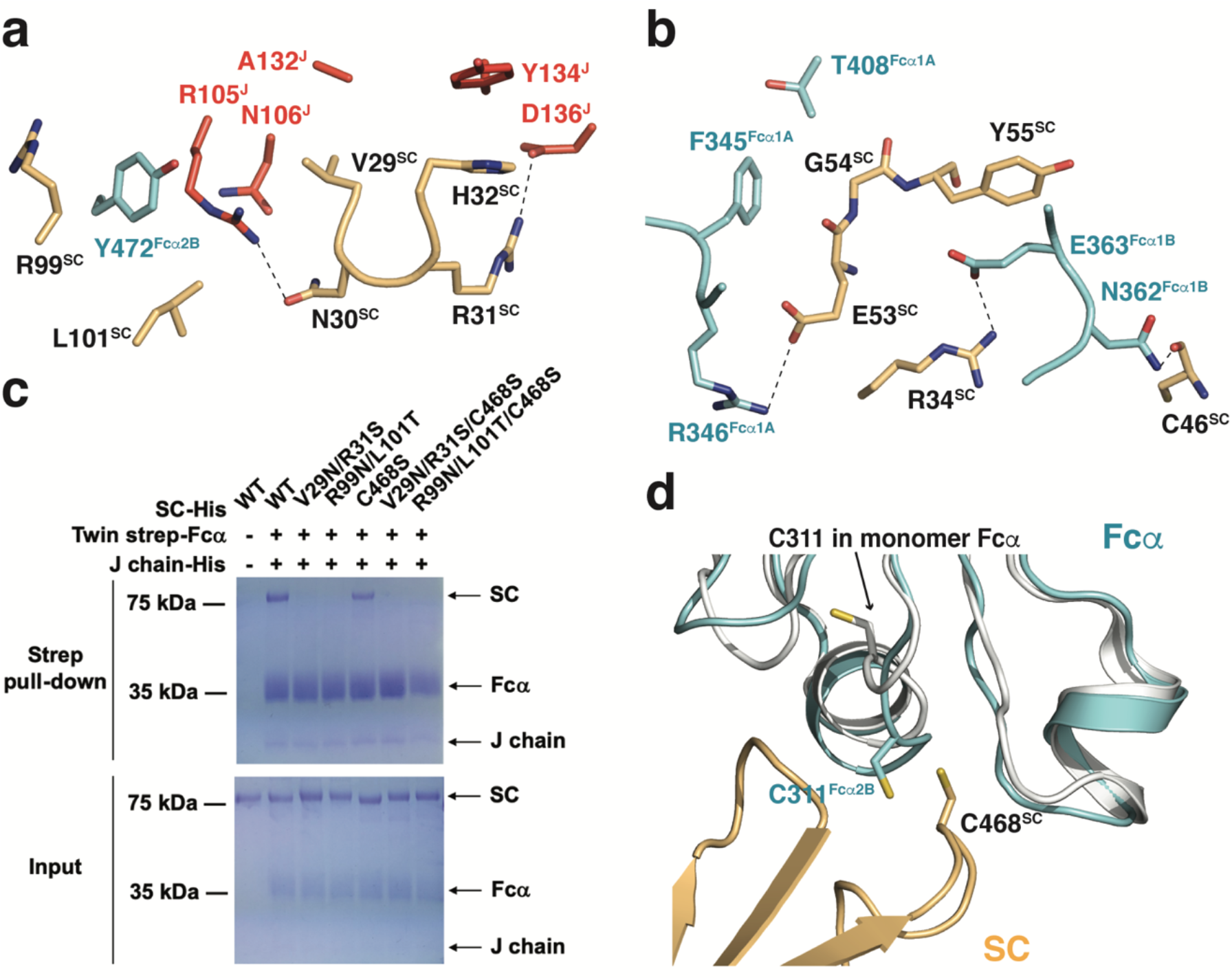
Interaction between dIgA and SC. **a.** Interactions between Fcα-J and SC at the CDR1 and CDR3 regions. Polar interactions are indicated by dashed lines. **b.** Direct interactions between Fcα and SC. **c.** SC mutants display reduced interactions with Fcα-J. **d.** The structure of the Fcα monomer (PDB ID: 2QEJ), shown in white, is overlaid onto Fcα2B in the Fcα-J-SC structure. Compared to Cys311 in the Fcα monomer, Cys311^Fcα2B^ flips out and can readily form a disulfide bond with Cys468^SC^.

The interaction between dIgA and pIgR/SC also uniquely involves the D5 domain of pIgR/SC, and a disulfide bond is formed between Cys468^SC^ and Cys311 in the Cα2 domain IgA ^23^. Indeed, although in the two determined structures of IgA ^29,30^, Cys311 is present in a hydrophobic pocket and not exposed, Cys311^Fcα2B^ flips out in the Fcα-J-SC complex and is located in close proximity to Cys468^SC^ (Fig. 4d). A disulfide bridge can be readily formed between them. Nevertheless, mutation of C468^SC^ only slightly decreased the binding between Fcα-J and SC in solution (Fig. 4c). This is consistent with previous analyses showing that the initial and primary association of dIgA with pIgR is mediated by interactions at the pIgR-D1 domain. Disulfide formation between dIgA and Cys468^SC^ is a late event during transcytosis, and is likely facilitated by the protein disulfide isomerases in secretory vesicles ^31^. The main function of this disulfide bond is to increase the stability of SIgA in the harsh environment of mucosal surfaces and external fluids.

### Interaction between SC and *S. pneumoniae* SpsA

SpsA comprises a C-terminal phosphorylcholine-binding domain that interacts with pneumococcal cell wall to function in bacterial colonization, and an N-terminal domain (NTD) that recruits host proteins including pIgR/SC ^15,32^. SpsA^NTD^ contains repeats of the leucine zipper motifs termed R1 and R2, each adopting a three-helix bundle structure ^33^. The YRNYPT hexapeptide motif involved in binding to pIgR/SC is located in the loop between helices α1 and α2 in the R1 motif. To reveal the molecular mechanism underlying the specific recognition of SIgA by SpsA, we reconstituted a Fcα-J-SC-SpsA^NTD^ quadruple complex and determined the cryo-EM structure at an overall resolution of 3.3 Å (Supplementary information, Figs. S1 and S4, Table S1). The α1-α2 loop of SpsA^NTD^, especially the YRNYPT hexapeptide motif, displays high-quality densities and can be clearly resolved (Supplementary information, Fig. S4b).

SpsA^NTD^ specifically interacts with the D3-D4 domains of human pIgR/SC ^34,35^. In the cryo-EM structure, the α1-α2 loop of SpsA^NTD^ docks into a pocket at the D3-D4 junction (Fig. 5a), formed by the DE loop of D3 and the C-C’ strands of D4. Notably, this pocket is only present in the ligand-bound conformation of SC ^17^. Tyr198, the first residue in the YRNYPT motif, forms a hydrogen bond with Tyr365^SC^ (Fig. 5b). Arg199 packs on Trp386^SC^, and form a salt bridge with Asp382^SC^. Asn200 forms a hydrogen bond with Arg376^SC^. Tyr201 packs on Pro283^SC^. Pro202 is surrounded by hydrophobic residues including Tyr365^SC^, Cys367^SC^, Cys377^SC^, Leu379^SC^, and Leu424^SC^. Substitution of Tyr201 with an Asp or Pro202 with a Glu abolished the binding of SpsA to SIgA ^36^. Thr203 interacts with Asn282^SC^. Notably, most of the SC residues described here are not conserved in pIgR/SC from other species (Supplementary information, Fig. S5), explaining the fact that SpsA only binds to human SIgA and pIgR/SC ^36^. Besides the interactions mediated by residues in the YRNYPT motif, Tyr206 forms a hydrogen bond with the main chain carbonyl group of Pro283^SC^, and Arg265 in helix α3 appears to hydrogen bond with the main chain carbonyl of Asp285^SC^ (Fig. 5b). These interactions further strengthen the binding between SpsA and SC.

**Figure 5.**
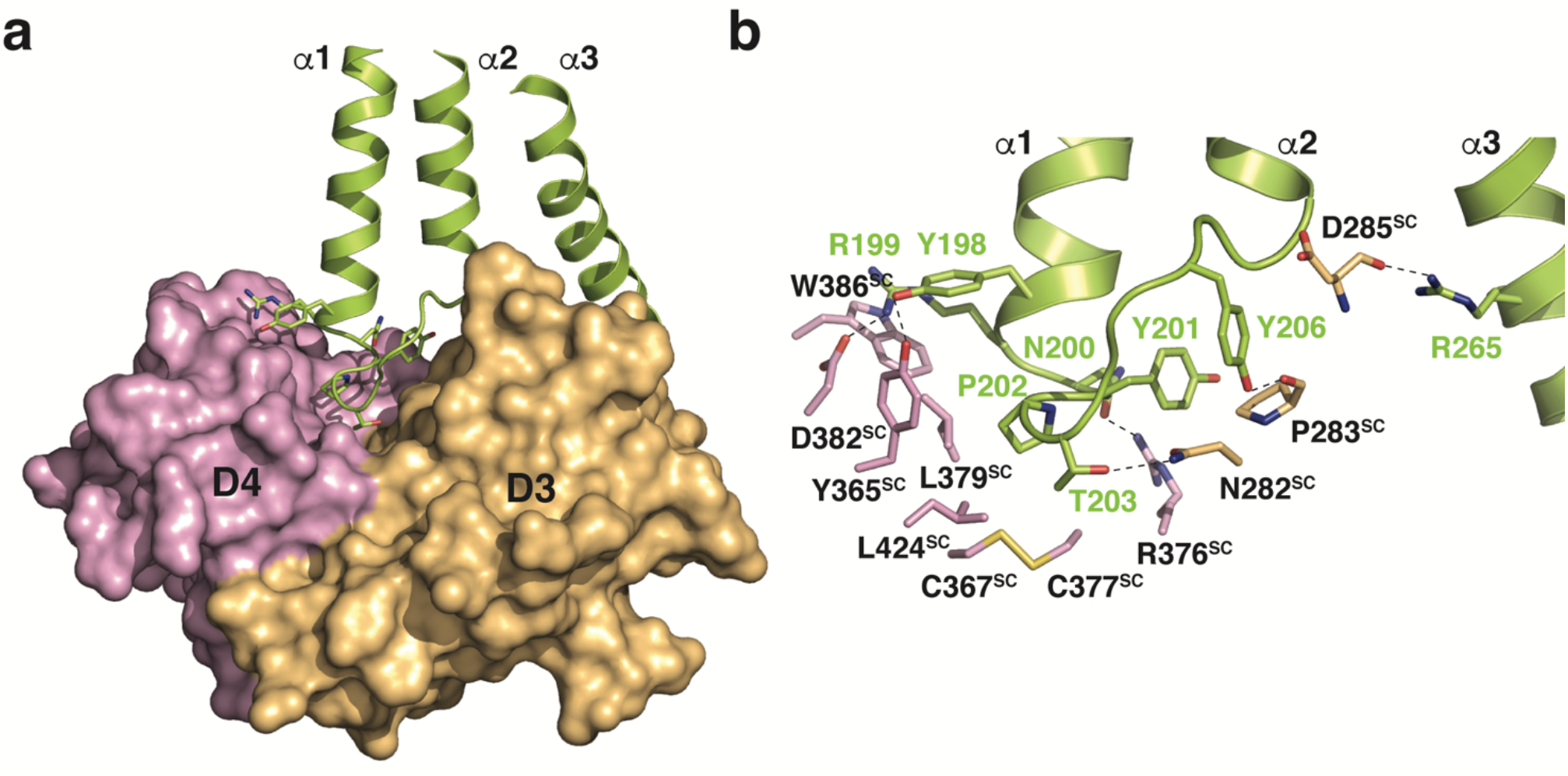
Interaction between SC and SpsA. **a.** Overall structure of the D3-D4 domains of SC in complex SpsA^NTD^. The D3-D4 domains of SC are shown in a surface representation, with D3 and D4 in gold and purple, respectively. SpsA^NTD^ is shown in lemon, and the side chains of the YRNYPT hexapeptide are depicted. **b.** Detailed interactions between SC and SpsA.

## Discussion

SIgA is of paramount importance to mucosal immunity. In adults, the daily synthesis of IgA is greater than all other types of antibody combined, and most of these IgA molecules are present in mucosal secretions in the form of dimeric SIgA. Despite the long history of SIgA research, its structure has remained elusive until only recently. During the preparation of this manuscript, the cryo-EM structures of SIgA have been published by Genentech ^37^. Our independent work reveals an architecture very similar to the dimeric SIgA core reported in this study, corroborating the reliability of these structures. IgA can induce immune signaling by binding to the IgA-specific receptor FcαRI/CD89 ^38,39^. It is not entirely clear whether monomeric IgA and SIgA can elicit similar immune responses. Crystal structure study reveals a 2:1 FcαRI:Fcα complex ^29^ (Fig. 6a). In secretory IgA, when the J-chain is present, only one side of the FcαRI-binding site would be exposed in each Fcα (Fig. 6b). The other side is occupied by the J-chain and not available for binding. From a structural point of view, there is no apparent reason to think that SIgA would not bind to FcαRI; nevertheless, it would have to bind membrane-bound FcαRI molecules in a different arrangement. Whether this altered mode of binding may account for the different immune responses elicited by monomeric IgA and SIgA remains to be investigated ^40^. Recently, human Fc receptor-like 3 (FCRL3) has been identified as a SIgA-specific receptor ^41^. It is likely that the J-chain and SC are involved in the interaction between SIgA and FCRL3, thereby contributing to the signaling function of SIgA.

**Figure 6.**
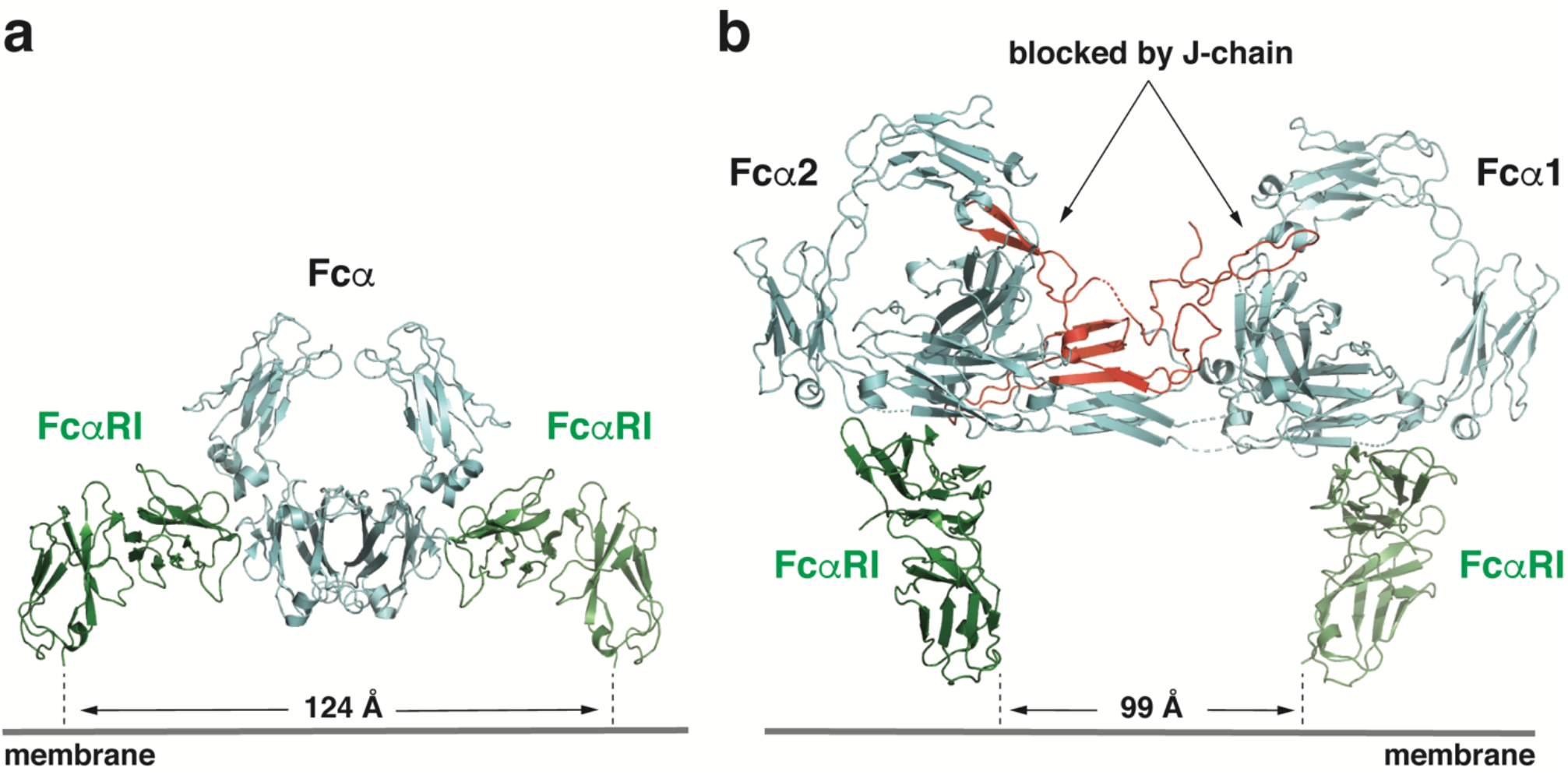
A hypothetical model of the dIgA-FcαRI complex. **a.** Crystal structure of the 2:1 FcαRI:Fcα complex (PDB ID: 1OW0). FcαRI is shown in green. The distance between the C-terminal ends of the two FcαRI molecules is indicated. **b.** In the Fcα-J structure, only one side of the FcαRI-binding site would be available. The other side is occupied by the J-chain and therefore not exposed for interacting with FcαRI.

*S. pneumoniae* is an important human pathogen. SpsA/CbpA/PspC is a major adhesin of *S. pneumoniae* and plays a role during its infection. Despite the fact that the DNA region encoding SpsA is highly polymorphic, the YRNYPT hexapeptide involved in binding to pIgR/SC is highly conserved, present in one or two copies in more than 70% strains of *S. pneumoniae* ^15,42^. In tissue culture models, SpsA-deficient *S. pneumoniae* showed greatly reduced ability to adhere to, and abolished activity to invade human cells ^16,32^. Notably, SpsA evolves to bind to human pIgR/SC specifically, since it does not interact with SIgA and SC from common laboratory animals including mouse, rat, rabbit, and guinea pig ^36^. Indeed, residues in pIgR/SC that participate in the interaction with SpsA are not conserved in these animals (Supplementary information, Fig. S5). These differences underscore the fact that *S. pneumoniae* is a human-specific pathogen and should be taken into consideration for the study of *S. pneumoniae* pathogenesis. On the other hand, the unique property of SpsA to bind human SC with high selectivity and affinity may allow the development of recombinant SpsA protein as a tool for efficient isolation and purification of human SIgA and SIgM.

## Materials and Methods

### Protein expression and purification

The DNA fragment encoding IgA1-Fc (residues 241-472) was cloned into a modified pcDNA vector with a N-terminal IL-2 signal peptide followed by a twin-strep tag. The DNA fragments encoding the full-length J-chain and SC were previously described ^17^. HEK293F cells were cultured in SMM 293T-I medium (Sino Biological Inc.) at 37 °C, with 5% CO_2_ and 55% humidity. The two plasmids expressing IgA1-Fc and J-chain were co-transfected into the cells using polyethylenimine (Polysciences). Four days after transfection, the conditioned media were collected by centrifugation, concentrated using a Hydrosart Ultrafilter (Sartorius), and exchanged into the binding buffer (25 mM Tris-HCl, pH 7.4, 150 mM NaCl). The recombinant proteins were isolated using Ni-NTA affinity purification and eluted with the binding buffer supplemented with 500 mM imidazole. The Fcα-J complex was further purified using a Superdex 200 increase column (GE Healthcare) and eluted using the binding buffer. SC was expressed and purified as previously described ^17^. To obtain the Fcα-J-SC tripartite complex, purified Fcα-J and SC were mixed in an 1:2 molar ratio and incubated on ice for 1 h. The complex was then further purified on a Superdex 200 increase column and eluted using the binding buffer.

The DNA fragment encoding SpsA^NTD^ (residues 38-324) was synthesized by Synbio Technologies and cloned into a modified pQlink vector with a N-terminal 8×His tag. SpsA^NTD^ was expressed in BL21(DE3)pLysS *E. coli*. The *E. coli* culture was grown in the Luria-Bertani medium at 37 °C to an OD_600_ of 0.8, and then induced with 0.5 mM isopropyl β-D-1-thiogalactopyranoside at 18 °C overnight for protein expression. The cells were collected by centrifugation, resuspended in the lysis buffer (50 mM Tris-HCl, pH 8.0, 300 mM NaCl, 1 mM phenylmethylsulfonyl fluoride), and then disrupted by sonication. The insoluble debris was removed by centrifugation. The recombinant protein was isolated using Ni-NTA affinity purification following standard procedure and eluted with 50 mM Tris-HCl, pH 8.0, 300 mM NaCl, and 500 mM imidazole. SpsA^NTD^ was then further purified by gel filtration chromatography using a Superdex 200 increase column (GE Healthcare) and eluted using the binding buffer. To obtain the Fcα-J-SC-SpsA^NTD^ quadruple complex, purified Fcα-J-SC and SpsA^NTD^ were mixed in an 1:2 molar ratio and incubated on ice for 1 h. The complex was then purified again on a Superdex 200 increase column and eluted using the binding buffer.

### Negative-staining and cryo-electron microscopy

The samples for EM study were prepared as previously described ^17^. All EM grids were evacuated for 2 minutes and glow-discharged for 30 seconds using a plasma cleaner (Harrick PDC-32G-2). For the negative-staining study, four-microliter aliquots of the Fcα-J-SC complex at 0.03 mg/ml were applied to glow-discharged carbon-coated copper grids (Zhong Jing Ke Yi, Beijing). After ~40 s, excessive liquid was removed using a filter paper (Whatman No. 1). The grid was then immediately stained using 2% uranyl acetate for 10 s and air dried. The grids were examined on a Tecnai G2 20 Twin electron microscope (FEI) operated at 120 kV. Images were recorded using a 4k × 4k CCD camera (Eagle, FEI). The Fcα-J-SC-SpsA^NTD^ sample was stained and examined similarly.

To prepare the sample for cryo-EM analyses, four-microliter aliquots of Fcα-J-SC (0.3 mg/ml) or Fcα-J-SC-SpsA^NTD^ (0.2 mg/ml) were applied to glow-discharged holy-carbon gold grids (Quantifoil, R1.2/1.3), blotted with filter paper at 4 °C and 100% humidity, and plunged into the liquid ethane using a Vitrobot Mark IV (FEI). Grids screening was performed using a Talos Arctica microscope equipped with Ceta camera (FEI). Data collection was carried out using a Titan Krios electron microscope (FEI) operated at 300 kV. Movies were recorded on a K2 Summit direct electron detector (Gatan) in a super resolution mode using the SerialEM software ^43^. A nominal magnification of 165,000X was used, and the exposure rate was 11.668 electrons per Å^2^ per second. The slit width of the energy filter was set to 20 eV. The defocus range was set from –0.8 to –1.6 μm. The micrographs were dose-fractioned into 32 frames with a total exposure time of 5.12 s and a total electron exposure of 60 electrons per Å^2^. Statistics for data collection are summarized in Supplementary information, Table S1.

### Imaging processing

For 3D reconstruction of the Fcα-J-SC complex, a total of 16,264 movie stacks were recorded. Raw movies frames were aligned and averaged into motion-corrected summed images with a pixel size of 0.828 Å by MotionCor2 ^44^. The contrast transfer function (CTF) parameters of each motion-corrected image were estimated by the Gctf program (v1.06) ^45^. Relion (v3.07) was used for all the following data processing ^46^. Manual screening was performed to remove low-quality images. A set of 475 particles were manually picked and subjected to 2D classification to generate templates for automatic particle picking. A total of 6,875,153 particles were then auto-picked, which were subjected to another round of 2D classification, resulting in 5,051,275 particles that were kept for the subsequent 3D classifications. Initial model was generated using Relion and used as a reference for 3D classification. Three of the six classes (665,589 particles) from the final round of 3D classification were selected and combined for refinement, resulting in a map with a 3.23 Å overall resolution after mask-based post-processing. Finally, Bayesian Polishing and CTF Refinement were applied, which yielded a density map at a resolution of 3.15 Å, based on the gold-standard FSC 0.143 criteria. The local resolution map was analyzed using ResMap ^47^ and displayed using UCSF Chimera ^48^. Similar data processing strategies were used for the Fcα-J-SC-SpsA complex. The workflows of data processing are illustrated in Supplementary information, Figure S1.

### Model building and structure refinement

The structure of Fcα (PDB ID: 1OW0), as well as the structures of the J-chain and SC from the Fcμ-J-SC complex (PDB ID: 6KXS), was docked into the EM map using Phenix ^49^ and then manually adjusted using Coot ^50^. The β5-β6 hairpin of the J-chain, which is disordered in the Fcμ-J-SC structure, was built de novo. The SpsA^NTD^ structure was also built de novo, using the previously determined solution structure of the R2 domain (PDB ID: 1W9R) as a reference. Residues in helices α1-α2 of SpsA^NTD^ can be unambiguously assigned. The amino acid registrations in helix α3 are not entirely reliable, since this helix is only loosely attached to SC and displays poor densities due to structural flexibility. Refinement was performed using the real-space refinement in Phenix. Figures were prepared with Pymol (Schrödinger) and UCSF Chimera.

### StrepTactin pull-down assay

WT and mutant SC proteins were purified using the Ni-NTA affinity method as previously described ^17^. For the pull-down experiments, they were first incubated with purified Fcα-J complex on ice for 1 h. The mixture was then incubated with the StrepTactin beads (Smart Lifesciences) in the binding buffer at 4 °C for another hour. A twin-strep tag is present on Fcα. The beads were spun down and then washed three times with the binding buffer. The bound proteins were eluted off the beads using the binding buffer supplemented with 10 mM desthiobiotin. The results were analyzed by SDS-PAGE and visualized by Coomassie staining.

## Acknowledgments

We thank the Core Facilities at the School of Life Sciences, Peking University for help with negative-staining EM; the Cryo-EM Platform of Peking University for help with data collection; the High-performance Computing Platform of Peking University for help with computation; the National Center for Protein Sciences at Peking University for assistance with Amersham Imager; and Guilan Li for help with cDNA library. The work was supported by the National Key Research and Development Program of China (2017YFA0505200, 2016YFC0906000 to J.X.; 2019YFA0508904 to N.G.), the National Science Foundation of China (31570735, 31822014 to J.X.; 31725007, 31630087 to N.G.), the Qidong-SLS Innovation Fund to J.X. and N.G.; and the Clinical Medicine Plus X Project of Peking University to J.X.

## Author contributions

Y.W. and Y.L. performed protein purification and biochemical experiments. Y.W. and G.W. prepared the cryo-EM sample and collected data. G.W. processed the cryo-EM data, under the supervision of N.G. J.X. built the structural model and wrote the manuscript, with inputs from all authors.

## Data availability

The cryo-EM maps and atomic coordinates of the Fcα-J-SC and Fcα-J-SC-SpsA complexes will be deposited in the EMDB and PDB, respectively.

---

**Figure S1.**
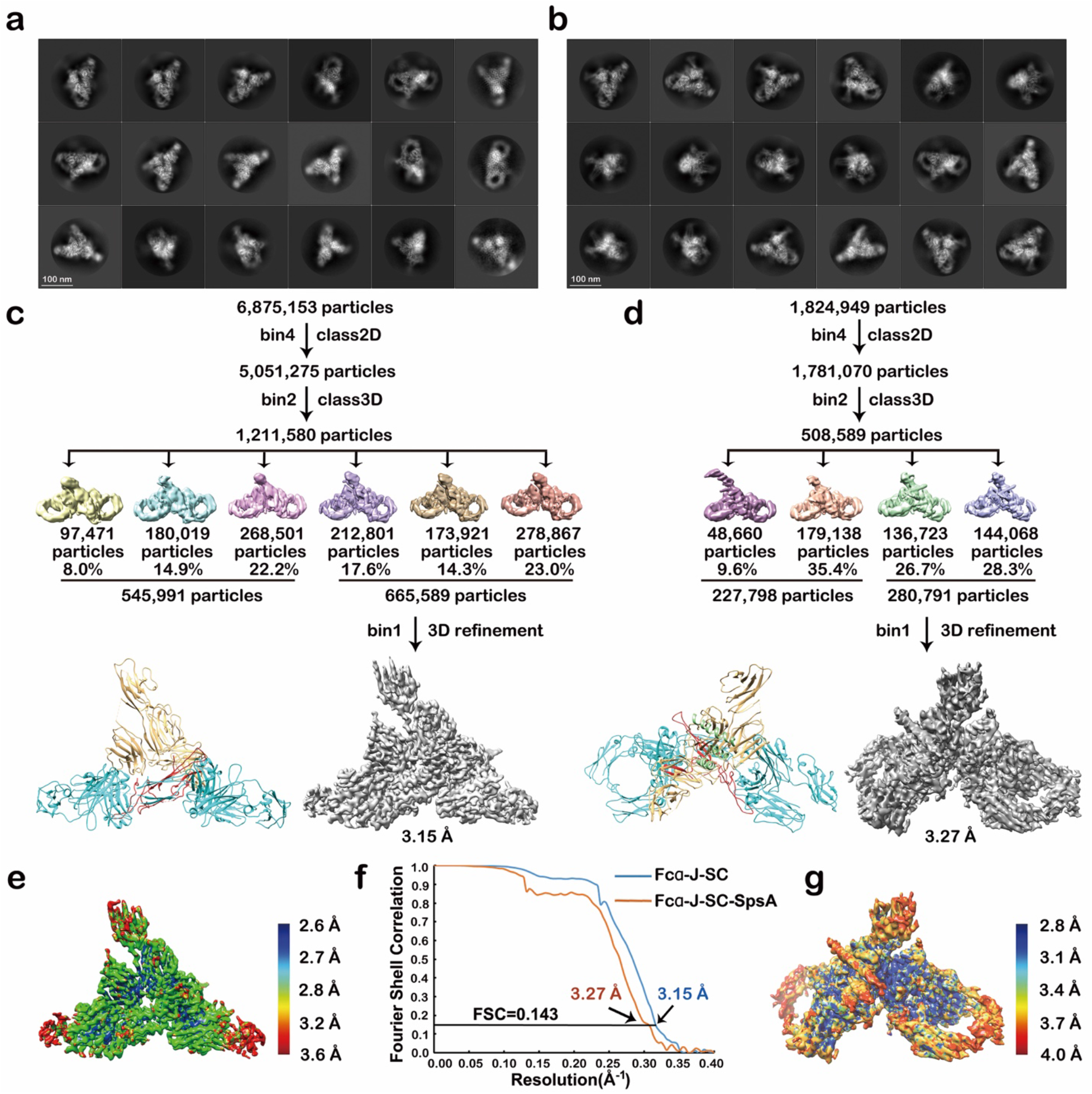
Workflows for cryo-EM 3D reconstructions. **a.** Representative 2D classes for the Fcα-J-SC complex. **b.** Representative 2D classes for the Fcα-J-SC-SpsA complex. **c.** Flow chart of data processing for the Fcα-J-SC complex. **d.** Flow chart of data processing for the Fcα-J-SC-SpsA complex. **e.** Local resolution estimation of the final map of Fcα-J-SC analyzed by ResMap. **f.** Gold standard Fourier shell correlation (FSC) curves with estimated resolutions. **g.** Local resolution estimation of the final map of Fcα-J-SC-SpsA.

**Figure S2.**
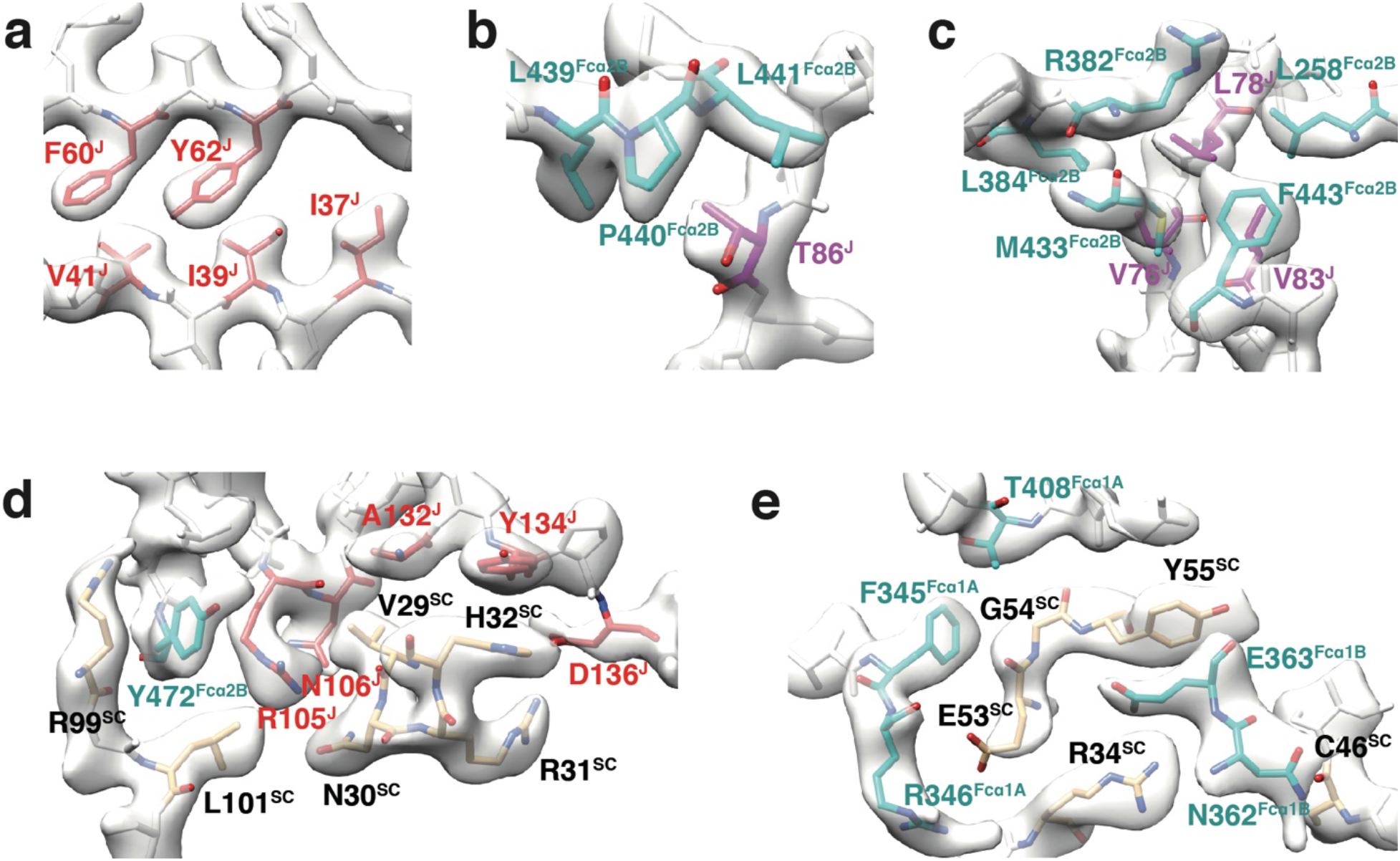
Density maps of selected regions in Fcα-J-SC.

**Figure S3.**
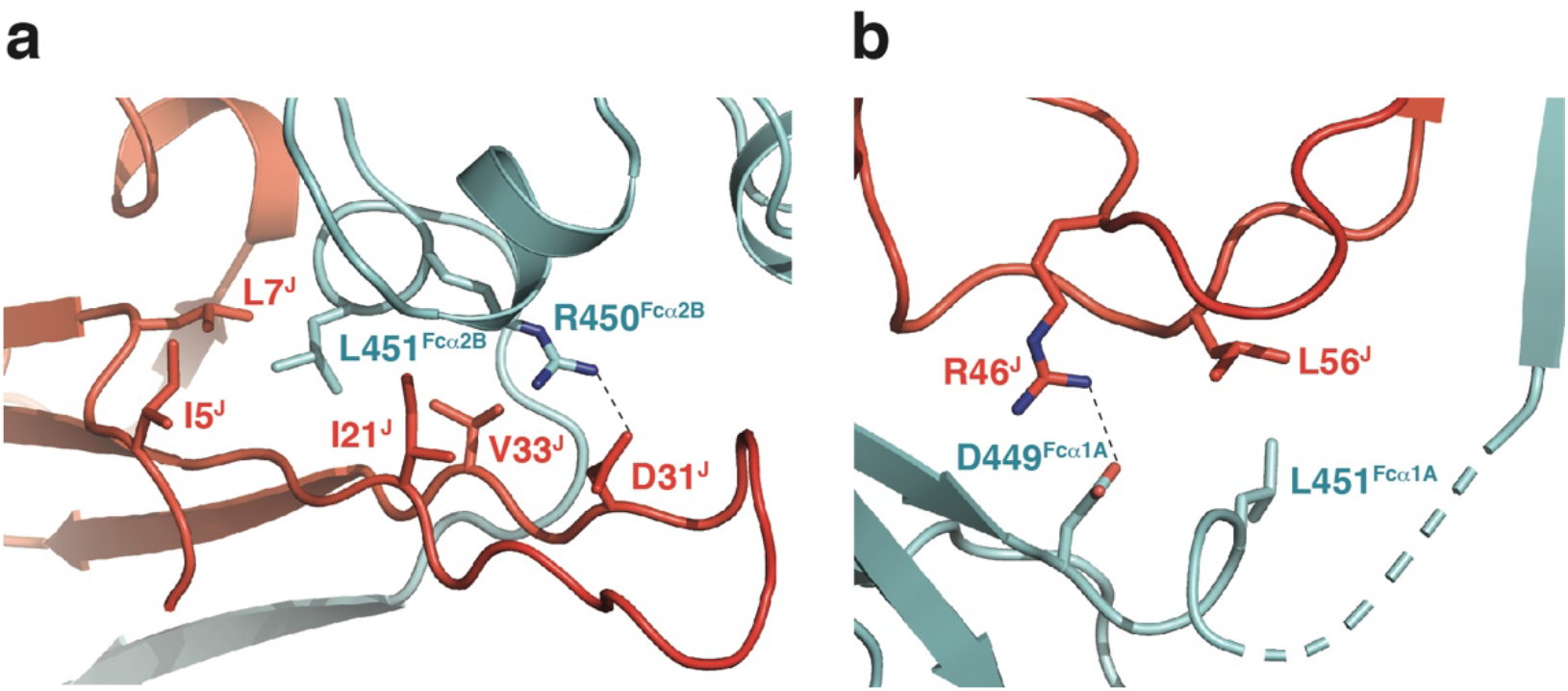
Interactions between the J-chain and Fcα. **a.** Interactions between the J-chain β2-β3 loop and the Fcα2B Cα3-tailpiece junction. **b.** Interactions between the J-chain β3-β4 loop and the Fcα1A Cα3-tailpiece junction.

**Figure S4.**
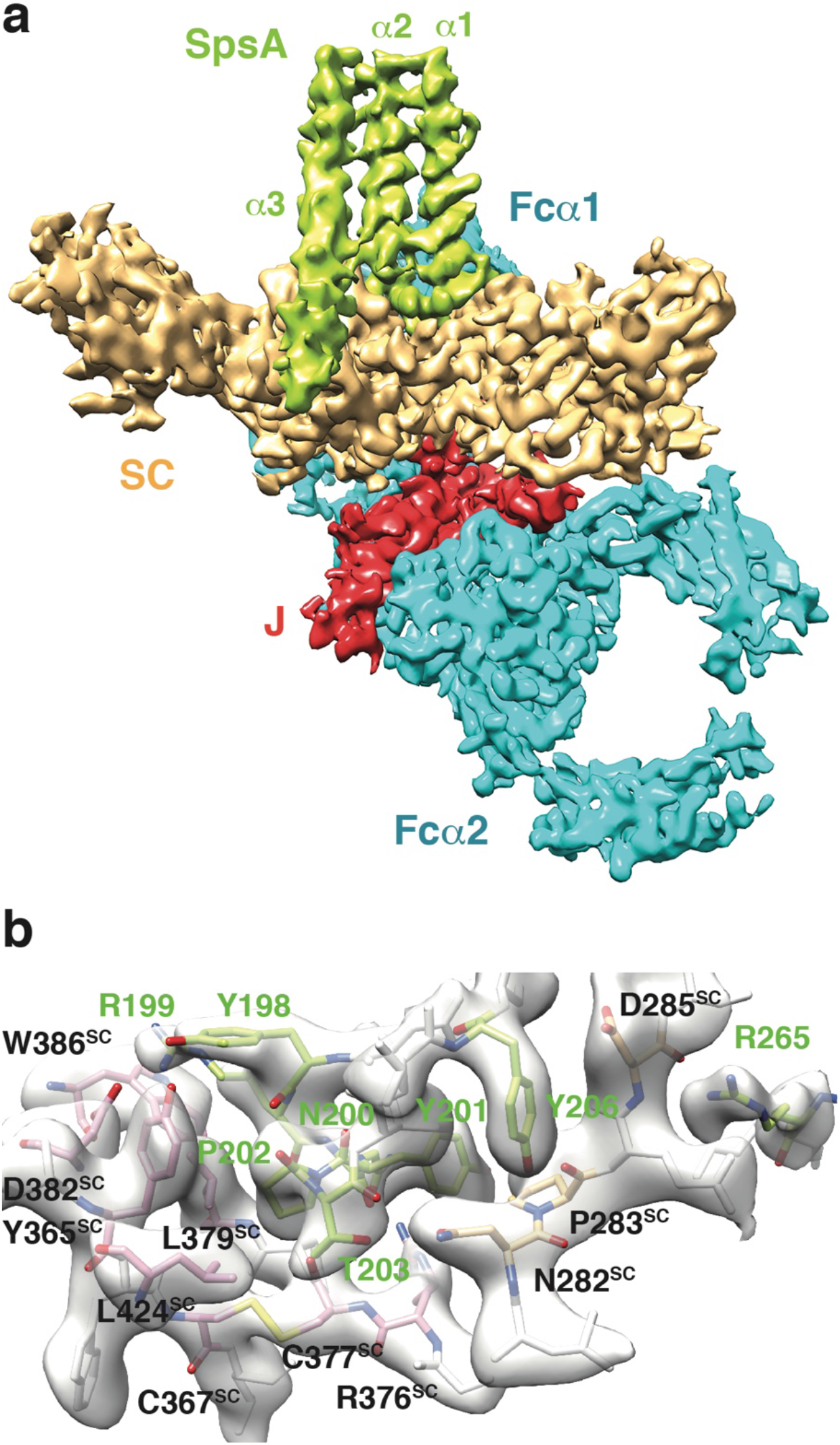
Cryo-EM structure of the Fcα-J-SC-SpsA complex. **a.** The cryo-EM density map of Fcα-J-SC-SpsA. **b.** Density maps of the SC-SpsA interface.

**Figure S5.**
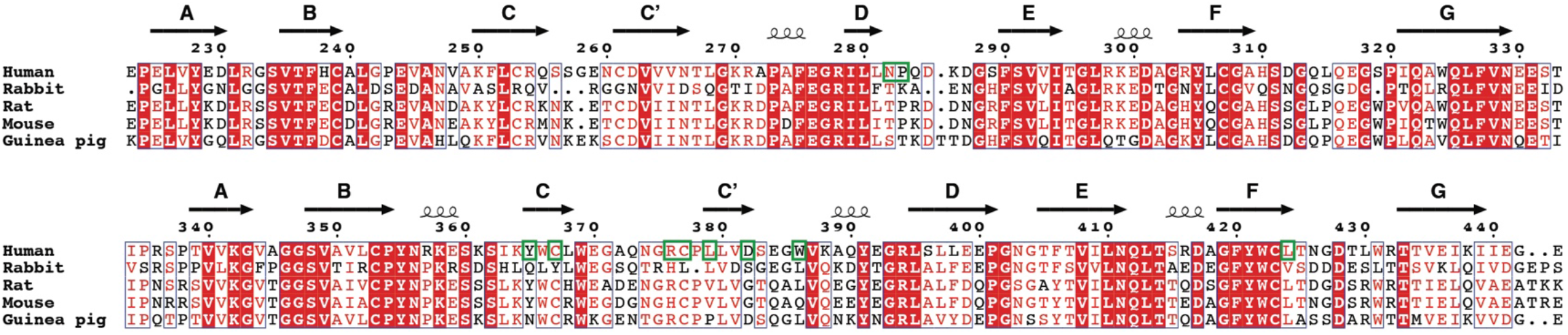
Sequence alignment of the D3-D4 domains of pIgR/SC from human and common laboratory animals. Residues in human pIgR/SC that are involved in binding to SpsA are highlighted in green boxes.

**Table S1.**
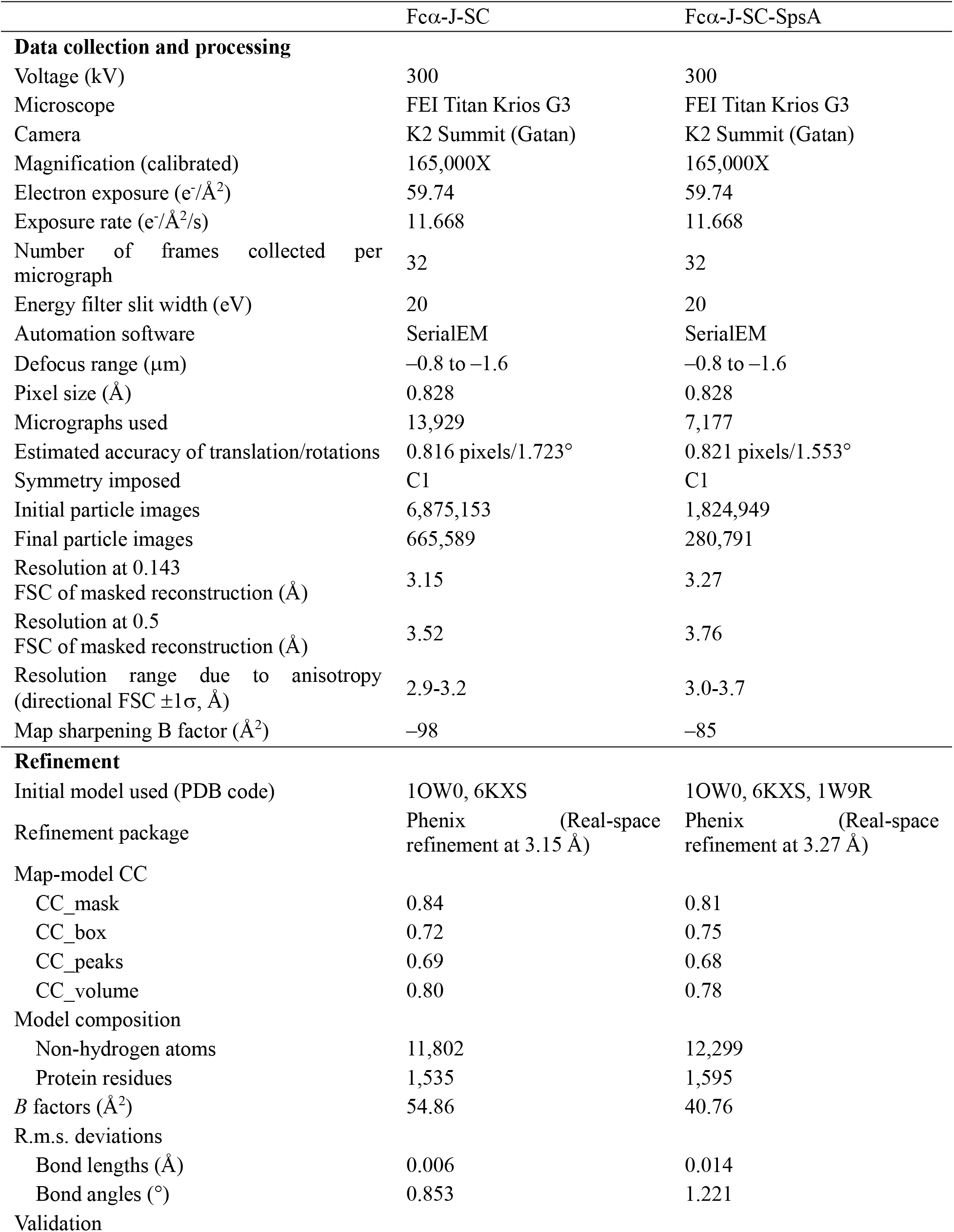
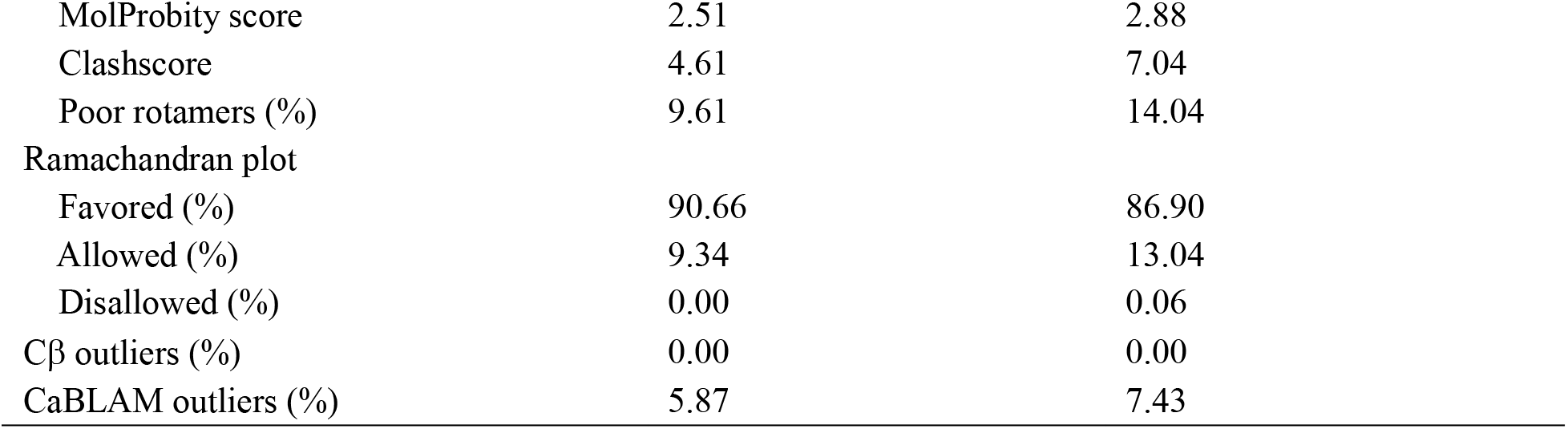
Cryo-EM data collection, processing and validation statistics

## Reference

1 Chodirker, W. B. & Tomasi, T. B., Jr. Gamma-Globulins: Quantitative Relationships in Human Serum and Nonvascular Fluids. Science 142, 1080–1081, doi:10.1126/science.142.3595.1080 (1963).

2 Halpern, M. S. & Koshland, M. E. Noval subunit in secretory IgA. Nature 228, 1276–1278, doi:10.1038/2281276a0 (1970).

3 Mestecky, J., Zikan, J. & Butler, W. T. Immunoglobulin M and secretory immunoglobulin A: presence of a common polypeptide chain different from light chains. Science 171, 1163–1165, doi:10.1126/science.171.3976.1163 (1971).

4 Koshland, M. E. The coming of age of the immunoglobulin J chain. Annu Rev Immunol 3, 425–453, doi:10.1146/annurev.iy.03.040185.002233 (1985).

5 Tomasi, T. B. The discovery of secretory IgA and the mucosal immune system. Immunol Today 13, 416–418, doi:10.1016/0167-5699(92)90093-M (1992).

6 Woof, J. M. & Mestecky, J. Mucosal immunoglobulins. Immunol Rev 206, 64–82, doi:10.1111/j.0105-2896.2005.00290.x (2005).

7 Mostov, K. E., Kraehenbuhl, J. P. & Blobel, G. Receptor-mediated transcellular transport of immunoglobulin: synthesis of secretory component as multiple and larger transmembrane forms. Proc Natl Acad Sci U S A 77, 7257–7261, doi:10.1073/pnas.77.12.7257 (1980).

8 Brandtzaeg, P. & Prydz, H. Direct evidence for an integrated function of J chain and secretory component in epithelial transport of immunoglobulins. Nature 311, 71–73, doi:10.1038/311071a0 (1984).

9 Norderhaug, I. N., Johansen, F. E., Schjerven, H. & Brandtzaeg, P. Regulation of the formation and external transport of secretory immunoglobulins. Crit Rev Immunol 19, 481–508 (1999).

10 Kaetzel, C. S. The polymeric immunoglobulin receptor: bridging innate and adaptive immune responses at mucosal surfaces. Immunol Rev 206, 83–99, doi:10.1111/j.0105-2896.2005.00278.x (2005).

11 Pabst, O., Cerovic, V. & Hornef, M. Secretory IgA in the Coordination of Establishment and Maintenance of the Microbiota. Trends Immunol 37, 287–296, doi:10.1016/j.it.2016.03.002 (2016).

12 Macpherson, A. J., Yilmaz, B., Limenitakis, J. P. & Ganal-Vonarburg, S. C. IgA Function in Relation to the Intestinal Microbiota. Annu Rev Immunol 36, 359–381, doi:10.1146/annurevimmunol-042617-053238 (2018).

13 Weiser, J. N., Ferreira, D. M. & Paton, J. C. Streptococcus pneumoniae: transmission, colonization and invasion. Nat Rev Microbiol 16, 355–367, doi:10.1038/s41579-018-0001-8 (2018).

14 Brooks, L. R. K. & Mias, G. I. Streptococcus pneumoniae’s Virulence and Host Immunity: Aging, Diagnostics, and Prevention. Front Immunol 9, 1366, doi:10.3389/fimmu.2018.01366 (2018).

15 Hammerschmidt, S., Talay, S. R., Brandtzaeg, P. & Chhatwal, G. S. SpsA, a novel pneumococcal surface protein with specific binding to secretory immunoglobulin A and secretory component. Mol Microbiol 25, 1113–1124, doi:10.1046/j.1365-2958.1997.5391899.x (1997).

16 Zhang, J. R. et al. The polymeric immunoglobulin receptor translocates pneumococci across human nasopharyngeal epithelial cells. Cell 102, 827–837, doi:10.1016/s0092-8674(00)00071-4 (2000).

17 Li, Y. et al. Structural insights into immunoglobulin M. Science, doi:10.1126/science.aaz5425 (2020).

18 Dourmashkin, R. R., Virella, G. & Parkhouse, R. M. Electron microscopy of human and mouse myeloma serum IgA. J Mol Biol 56, 207–208, doi:10.1016/0022-2836(71)90097-0 (1971).

19 Bonner, A., Furtado, P. B., Almogren, A., Kerr, M. A. & Perkins, S. J. Implications of the near-planar solution structure of human myeloma dimeric IgA1 for mucosal immunity and IgA nephropathy. J Immunol 180, 1008–1018, doi:10.4049/jimmunol.180.2.1008 (2008).

20 Bastian, A., Kratzin, H., Eckart, K. & Hilschmann, N. Intra- and interchain disulfide bridges of the human J chain in secretory immunoglobulin A. Biol Chem Hoppe Seyler 373, 1255–1263, doi:10.1515/bchm3.1992.373.2.1255 (1992).

21 Frutiger, S., Hughes, G. J., Paquet, N., Luthy, R. & Jaton, J. C. Disulfide bond assignment in human J chain and its covalent pairing with immunoglobulin M. Biochemistry 31, 12643–12647, doi:10.1021/bi00165a014 (1992).

22 Frutiger, S., Hughes, G. J., Hanly, W. C., Kingzette, M. & Jaton, J. C. The amino-terminal domain of rabbit secretory component is responsible for noncovalent binding to immunoglobulin A dimers. J Biol Chem 261, 16673–16681 (1986).

23 Fallgreen-Gebauer, E. et al. The covalent linkage of secretory component to IgA. Structure of sIgA. Biol Chem Hoppe Seyler 374, 1023–1028 (1993).

24 Coyne, R. S., Siebrecht, M., Peitsch, M. C. & Casanova, J. E. Mutational analysis of polymeric immunoglobulin receptor/ligand interactions. Evidence for the involvement of multiple complementarity determining region (CDR)-like loops in receptor domain I. J Biol Chem 269, 31620–31625 (1994).

25 Hexham, J. M. et al. A human immunoglobulin (Ig)A calpha3 domain motif directs polymeric Ig receptor-mediated secretion. J Exp Med 189, 747–752, doi:10.1084/jem.189.4.747 (1999).

26 Hamburger, A. E., West, A. P., Jr. & Bjorkman, P. J. Crystal structure of a polymeric immunoglobulin binding fragment of the human polymeric immunoglobulin receptor. Structure 12, 1925–1935, doi:10.1016/j.str.2004.09.006 (2004).

27 Lewis, M. J., Pleass, R. J., Batten, M. R., Atkin, J. D. & Woof, J. M. Structural requirements for the interaction of human IgA with the human polymeric Ig receptor. J Immunol 175, 6694–6701, doi:10.4049/jimmunol.175.10.6694 (2005).

28 Stadtmueller, B. M. et al. The structure and dynamics of secretory component and its interactions with polymeric immunoglobulins. Elife 5, doi:10.7554/eLife.10640 (2016).

29 Herr, A. B., Ballister, E. R. & Bjorkman, P. J. Insights into IgA-mediated immune responses from the crystal structures of human FcalphaRI and its complex with IgA1-Fc. Nature 423, 614–620, doi:10.1038/nature01685 (2003).

30 Ramsland, P. A. et al. Structural basis for evasion of IgA immunity by Staphylococcus aureus revealed in the complex of SSL7 with Fc of human IgA1. Proc Natl Acad Sci U S A 104, 15051–15056, doi:10.1073/pnas.0706028104 (2007).

31 Chintalacharuvu, K. R. et al. Disulfide bond formation between dimeric immunoglobulin A and the polymeric immunoglobulin receptor during hepatic transcytosis. Hepatology 19, 162–173 (1994).

32 Rosenow, C. et al. Contribution of novel choline-binding proteins to adherence, colonization and immunogenicity of Streptococcus pneumoniae. Mol Microbiol 25, 819–829, doi:10.1111/j.1365-2958.1997.mmi494.x (1997).

33 Luo, R. et al. Solution structure of choline binding protein A, the major adhesin of Streptococcus pneumoniae. EMBO J 24, 34–43, doi:10.1038/sj.emboj.7600490 (2005).

34 Lu, L., Lamm, M. E., Li, H., Corthesy, B. & Zhang, J. R. The human polymeric immunoglobulin receptor binds to Streptococcus pneumoniae via domains 3 and 4. J Biol Chem 278, 48178–48187, doi:10.1074/jbc.M306906200 (2003).

35 Elm, C. et al. Ectodomains 3 and 4 of human polymeric Immunoglobulin receptor (hpIgR) mediate invasion of Streptococcus pneumoniae into the epithelium. J Biol Chem 279, 6296–6304, doi:10.1074/jbc.M310528200 (2004).

36 Hammerschmidt, S., Tillig, M. P., Wolff, S., Vaerman, J. P. & Chhatwal, G. S. Species-specific binding of human secretory component to SpsA protein of Streptococcus pneumoniae via a hexapeptide motif. Mol Microbiol 36, 726–736, doi:10.1046/j.1365-2958.2000.01897.x (2000).

37 Kumar, N., Arthur, C. P., Ciferri, C. & Matsumoto, M. L. Structure of the secretory immunoglobulin A core. Science, doi:10.1126/science.aaz5807 (2020).

38 van Egmond, M. et al. IgA and the IgA Fc receptor. Trends Immunol 22, 205–211, doi:10.1016/s1471-4906(01)01873-7 (2001).

39 Monteiro, R. C. & Van De Winkel, J. G. IgA Fc receptors. Annu Rev Immunol 21, 177–204, doi:10.1146/annurev.immunol.21.120601.141011 (2003).

40 van Egmond, M. et al. FcalphaRI-positive liver Kupffer cells: reappraisal of the function of immunoglobulin A in immunity. Nat Med 6, 680–685, doi:10.1038/76261 (2000).

41 Agarwal, S. et al. Human Fc Receptor-like 3 Inhibits Regulatory T Cell Function and Binds Secretory IgA. Cell Rep 30, 1292–1299 e1293, doi:10.1016/j.celrep.2019.12.099 (2020).

42 Iannelli, F., Oggioni, M. R. & Pozzi, G. Allelic variation in the highly polymorphic locus pspC of Streptococcus pneumoniae. Gene 284, 63–71, doi:10.1016/s0378-1119(01)00896-4 (2002).

43 Mastronarde, D. N. Automated electron microscope tomography using robust prediction of specimen movements. J Struct Biol 152, 36–51, doi:10.1016/j.jsb.2005.07.007 (2005).

44 Zheng, S. Q. et al. MotionCor2: anisotropic correction of beam-induced motion for improved cryoelectron microscopy. Nat Methods 14, 331–332, doi:10.1038/nmeth.4193 (2017).

45 Zhang, K. Gctf: Real-time CTF determination and correction. J Struct Biol 193, 1–12, doi:10.1016/j.jsb.2015.11.003 (2016).

46 Zivanov, J. et al. New tools for automated high-resolution cryo-EM structure determination in RELION-3. Elife 7, doi:10.7554/eLife.42166 (2018).

47 Kucukelbir, A., Sigworth, F. J. & Tagare, H. D. Quantifying the local resolution of cryo-EM density maps. Nat Methods 11, 63–65, doi:10.1038/nmeth.2727 (2014).

48 Pettersen, E. F. et al. UCSF Chimera—A Visualization System for Exploratory Research and Analysis. J Comput Chem 25, 1605–1612, doi:10.1002/jcc.20084 (2004).

49 Adams, P. D. et al. PHENIX: A comprehensive Python-based system for macromolecular structure solution. Acta Crystallographica Section D: Biological Crystallography 66, 213–221, doi:10.1107/S0907444909052925 (2010).

50 Emsley, P., Lohkamp, B., Scott, W. G. & Cowtan, K. Features and development of Coot. Acta Crystallographica Section D: Biological Crystallography 66, 486–501, doi:10.1107/S0907444910007493 (2010).

